# Regulation of eIF4F complex by the peptidyl prolyl isomerase FKBP7 in taxane-resistant prostate cancer

**DOI:** 10.1101/096271

**Authors:** Marine F. Garrido, Nicolas J-P. Martin, Catherine Gaudin, Frédéric Commo, Nader AL Nakouzi, Ladan Fazli, Elaine Del Nery, Jacques Camonis, Franck Perez, Stéphanie Lerondel, Alain LE Pape, Hussein Abou-Hamdan, Martin Gleave, Yohann Loriot, Laurent Désaubry, Stephan Vagner, Karim Fizazi, Anne Chauchereau

## Abstract

Targeted therapies that exploit the signaling pathways involved in prostate cancer are required to overcome chemoresistance and improve treatment outcomes for men. Molecular chaperones play a key role in the regulation of protein homeostasis and are potential targets to alleviate chemoresistance. Using image-based high content siRNA functional screening based on a gene expression signature, we identified FKBP7, a molecular chaperone overexpressed in docetaxel-resistant and in cabazitaxel-resistant prostate cancer cells. FKBP7 was upregulated in human prostate cancers and correlated with the recurrence in patients receiving Docetaxel. *FKBP7* silencing showed that FKBP7 is required to maintain the growth of chemoresistant cell lines and of chemoresistant tumors in mice. Mass spectrometry analysis revealed that FKBP7 interacts with the eIF4G component of the eIF4F translation initiation complex to mediate survival of chemoresistant cells. Using small molecule inhibitors of eIF4A, the RNA helicase component of eIF4F, we were able to overcome docetaxel and cabazitaxel resistance.

## INTRODUCTION

Molecular chaperones are upregulated and associated with treatment resistance in castration-resistant prostate cancer (CRPC) (Ischia *et al*, 2013; Lamoureux *et al*, 2014), an advanced form of prostate cancer that develops when the disease progresses following initial treatment with surgery and/or medical castration with androgen deprivation therapy (ADT). Docetaxel and cabazitaxel are two taxane chemotherapies that are approved for treatment of metastatic CRPC (Hurwitz, 2015). Recent studies have reported benefits when docetaxel was combined with ADT during the hormone-sensitive stage of the disease (Hurwitz, 2015; James *et al*, 2016). Approximately 50% of patients do not respond to docetaxel. Mechanisms of resistance to docetaxel include decreased cellular accumulation of the drug due to overexpression of membrane-bound efflux proteins, expression of tubulin isotopes, and defects in apoptosis (Seruga *et al*, 2011; Mahon *et al*, 2011). Many of these pathways involve molecular chaperones, but targeting these proteins have previously been developed to combat docetaxel resistance with only limited efficacy reported to date. New therapeutic strategies are needed to overcome taxane-resistance and improve patient outcomes for men with prostate cancer.

Several molecular chaperones, including the heat-shock proteins Hsp27 and Hsp90, clusterin (CLU), and the FKB506-binding proteins (FKBPs) are involved in protein folding, cellular signaling, apoptosis and transcription, and are potential targets for cancer treatment (Solassol *et al*, 2011; Zoubeidi & Gleave, 2012). FKBP12 (*FKBP1A*) was the first enzyme discovered to bind FK506, a natural immunosuppressant and rapamycin, a macrolide indicated in prophylaxis of organ rejection in renal transplantation (Bierer *et al*, 1990). The FKBP12-rapamycin complex associates with the major downstream Akt target mammalian target of rapamycin (mTOR)-kinase, and has both immunosuppressive and antiproliferative properties. Larger FKBPs, such as FKBP52 (*FKBP4*) and FKBP51 (*FKBP5*), have also been shown to form rapamycin-induced ternary complexes that inhibit mTOR-kinase activity. FKBP51 and FKBP52 regulate the microtubule-associated protein tau and thus affect microtubule stability (Cioffi *et al*, 2011). FKBP7, also known as FKBP23, is another molecular chaperone that was previously cloned from mouse heart (Nakamura *et al*, 1998). FKBP7 is located in the endoplasmic reticulum and was shown to suppress the ATPase activity of the mouse ER chaperone HSPA5/GRP78/Bip by its prolyl-peptidyl isomerase activity (Zhang *et al*, 2004; Nakamura *et al*, 1998). The function of the human FKBP7 has not yet been characterized and to date its role in cancer has never been explored.

In the present study, we established four chemoresistant prostate cancer cell lines to explore mechanisms of taxane resistance. A high throughput proteomic approach was used to investigate the signaling pathway involving FKBP7 and its potential role in taxane resistance.

## RESULTS

### FKBP7 is upregulated during the progression of chemoresistant CRPC

To decipher the mechanisms of taxane resistance in prostate cancer, we developed a series of four isogenic parental, docetaxel-resistant (Dtx-R), and cabazitaxel-resistant (Cbx-R) cell lines representative of the different types of epithelial prostate cancer cells (Fig S1). By comparing the gene expression profiles between parental and different docetaxel-resistant cells (Chauchereau *et al*, 2011; Nakouzi *et al*, 2014), we generated a signature of 998 highly differentially expressed genes potentially correlated with chemoresistance (Table S1). Following image-based high-content screening in which the 593 overexpressed genes were independently targeted with 4 different siRNAs in IGR-CaP1-R100 cells, we identified 34 genes required for cell survival of docetaxel-resistant cells, for which at least 2 siRNAs showed robust Z-score>2 for G0 cell-cycle arrest phenotype modification and cell proliferation. We focused our attention on chaperone proteins responsive to stress. In particular, as already reported in several solid cancers (Gifford *et al*, 2016; Roller & Maddalo, 2013), the ER chaperone protein HSPA5 also known as GRP78/BiP, was sorted as a candidate involved in chemoresistance in our model. Considering that HSPA5 was reported to interact with the FKBP7 chaperone in a mouse model (Zhang *et al*, 2004), we focused on the potential role of the uncharacterized human protein FKBP7 in the mechanism of taxane resistance in prostate cancer.

FKBP7 protein levels were higher in the four parental prostate cancer cell lines compared to the non-cancerous prostate cells RWPE-1 (Fig 1A). FKBP7 protein levels were higher in all docetaxel- and cabazitaxel-resistant cells compared to their respective parental cells, with an 8 fold-change in taxane-resistant IGR-CaP1 cells. Increased FKBP7 protein level is related to gene expression upregulation since we found more FKBP7 mRNA in the taxane-resistant cells (Fig S2). To investigate the physiological relevance of FKBP7 expression in prostate cancer, we determined FKBP7 expression by immunohistochemistry using tissue microarrays containing 381 prostate cancers and benign tissues obtained from radical prostatectomy or from transurethral resection (Table S2).

**Figure-1.**
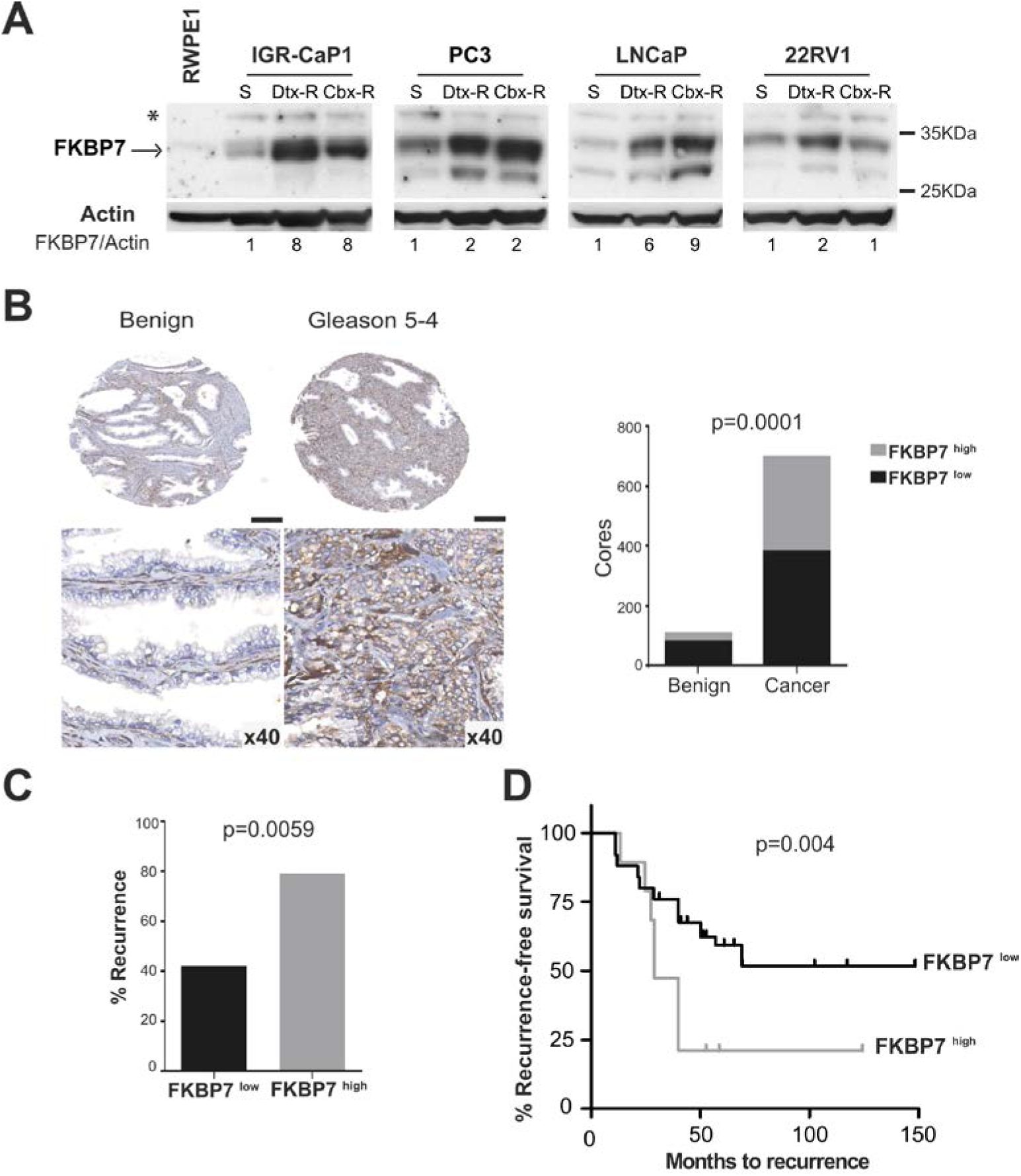
FKBP7 is upregulated during the progression of chemoresistant CRPC. (A) Immunoblot of FKBP7 protein expression in the non-cancerous prostate cells RWPE-1, the parental (S) IGR-CaP1, PC3, LNCaP and 22RV1, the docetaxel-resistant cells (Dtx-R) and the cabazitaxel-resistant cells (Cbx-R). Actin is the loading control. FKBP7 protein level has been quantified with the Image Lab software: FKBP7 in resistant cells is expressed relatively to their parental cell lines. *indicates non-specific band. (B) On left panel, representative immunohistochemistry images of FKBP7 staining in benign and cancer prostate tissues. Scale bars: 200μm (top). On right panel, quantification of FKBP7 protein level in benign prostate tissue and tumor. Chi-square test p=0.0001. FKBP7 low corresponds to a score of 0 and 1; FKBP7 high corresponds to a score of 2 and 3. n=808. (C) Correlation of FKBP7 expression intensity with the percentage of recurrence in 69 patients treated with docetaxel. Fisher’s exact test p=0.0059. Event is defined as any recurrence or metastasis or prostate cancer death post-surgery. The date of surgery was set as baseline. (D) Kaplan-Meir plot representing the recurrence-free survival (RFS) associated with the FKBP7 staining in TMA obtained from 69 patients who received docetaxel as neoadjuvant therapy. The association between time to recurrence (months) and staining status of FKBP7 (high or low), where an event is defined as PSA recurrence, metastasis or prostate cancer death, was calculated with Cox proportional hazard model: Hazard ratio 0.3846 (95% confidence interval [CI]: 0.195 to 0.7586); p=0.004 (log rank test).

Consistent with what we observed in cell lines, FKBP7 was more present in different stages of prostate cancers compared to benign tissues (Fig 1B). Quantification of staining intensities showed that FKBP7 levels (moderate to strong staining) were significantly higher in prostate cancer than in benign tissues (Fig 1B). We further evaluated FKBP7 expression using tissue microarrays comprising a subset of 69 prostate cancers from patients who received docetaxel as neoadjuvant therapy. The levels of FKBP7 significantly correlated with the percentage of recurrence in patients (defined as any events of recurrence, metastasis or prostate cancer death post-surgery): 79% of patients with high FKBP7 levels developed recurrence compared to 42% for patients with a low level of FKBP7. High levels of FKBP7 were significantly associated with a shorter time to recurrence in patients (p=0.004, hazard ratio for low FKBP7 staining: 0.3846) (Fig 1D). The levels of FKBP7 during the progression to lethal prostate cancer did not correspond to gene mutations in the published data from whole exome sequencing in docetaxel-treated CRPC patients (Grasso *et al*, 2012). Together, these results provide strong evidence for a role of FKBP7 in prostate tumor cell growth and chemoresistance.

### SiRNA-mediated knock-down of FKBP7 blocks chemoresistant cell growth and sensitizes chemoresistant cells to taxane treatment

To determine whether FKBP7 could be a therapeutic target in chemoresistant prostate cancer, we used two different siRNA sequences targeting FKBP7, to knock-down FKBP7 expression in all chemoresistant cells. FKBP7 silencing significantly decreased chemoresistant cell growth (Fig 2A). FKBP7 silencing slightly induced apoptosis of chemoresistant cells and PARP cleavage was further increased after treatment with docetaxel and cabazitaxel (Fig 2B). These results show the importance of FKBP7 signaling in taxane-acquired chemoresistance.

**Figure 2.**
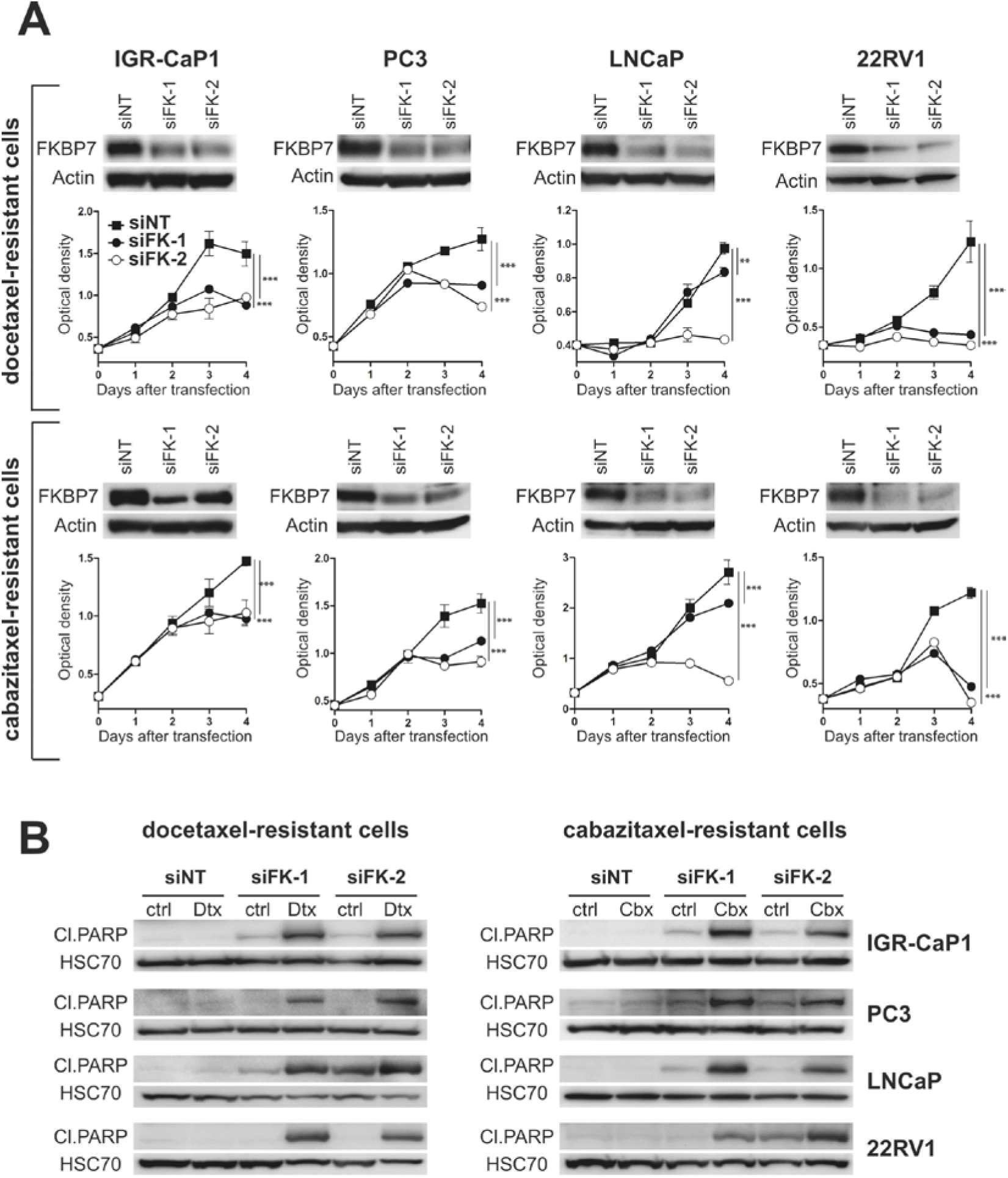
SiRNA-mediated knock-down of FKBP7 blocks chemoresistant cell growth and sensitizes chemoresistant cells to taxane treatment. (**A**) Immunoblot shows FKBP7 knockdown efficiency 48h after transfection with either two different siRNA sequences targeting FKBP7 (siFK-1 or si-FK-2) or control siRNA (siNT). Actin is the loading control. Cell viability with cells transfected with either two different siRNA sequences targeting FKBP7 or control siRNA is determined daily with WST1 assay. Data represent the mean ± SD (n=3). **p<0.01;***p<0.001 as determined by two-way ANOVA with Bonferroni posttests. Experiments were performed with either docetaxel-resistant cells (top panel) or cabazitaxel-resistant cells (bottom). (**B**) Immunoblots showing cleaved (Cl.) PARP protein in either docetaxel-resistant cells (left panel) or cabazitaxel-resistant cells (right panel) after 96h of transfection with siRNA ctrl (siNT) or siFKBP7s (siFK-1 or siFK-2) alone or combined with docetaxel or cabazitaxel treatment for 72h. Chemoresistant cells were treated with their respective maximal dose of resistance (Supplemental Figure 1). HSC70 is the loading control.

### FKBP7 silencing reduces tumor growth in a docetaxel-resistant mouse model

The IGR-CaP1 cell line was used to generate a chemoresistant mCRPC mouse model. Various clones of IGR-CaP1 resistant to different doses of docetaxel were injected subcutaneously into nude mice. The 25nM-resistant emerging tumor was maintained *in vivo* for 5 successive passages to increase the tumorigenicity (Fig S3). These tumors, called IGR-CaP1-Rvivo, did not respond to 3 successive injections of docetaxel at 30mg/kg, in contrast to the parental IGR-CaP1 mouse model which responded to docetaxel (Fig 3A), and therefore constitute a new docetaxel-resistant mouse model. A new cell line, called IGR-CaP1-Rvivo, was generated from the IGR-CaP1-Rvivo tumors and showed chemoresistance characteristics (with IC_50_=207nM towards docetaxel) (Fig 3B, left panel). In addition, and in agreement with our results obtained *in vitro* and in human samples, the resistant IGR-CaP1-Rvivo cell line exhibited high levels of FKBP7 (Fig 3B, right panel). Thus, to interrogate whether FKBP7 could be a therapeutic target in a docetaxel-resistant mouse model, we established two IGR-CaP1-Rvivo cell lines in which FKBP7 was stably silenced with two different shRNAs. We achieved a 90% (with shFK-a) and 40% (with shFK-b) reduction of FKBP7 levels in cells stably expressing the shRNAs targeting FKBP7, compared to control shRNA expressing cells (Fig 3C). ShRNA-transduced cell lines were subsequently injected subcutaneously in immunodeficient mice. In the absence of docetaxel, tumor growth was significantly reduced with FKBP7 depletion (68% and 47% inhibition of tumor growth in shFK-a- and shFK-b-transduced cells, respectively), compared to the control shRNA-transduced cell line (Fig 3C, left panel). This effect was more pronounced in the xenograft issued from the shFK7-a cells exhibiting less FKBP7. This suggests that FKBP7 sustained the growth of chemoresistant tumors. Treatment of mice with docetaxel, when tumors reached 450-500mm3, strongly abrogated the growth of shFK-a- transduced tumors (exhibiting the lowest level of FKBP7), while it weakly decreased the growth of control shRNA-transduced tumors (Fig 3C, right panel). Thus, our results demonstrated that both *in vitro* and *in vivo*, FKBP7 silencing inhibits cell proliferation and sensitizes chemoresistant cells to taxanes, showing that FKBP7 can be a relevant therapeutic target to overcome chemoresistance in prostate cancer.

**Figure 3.**
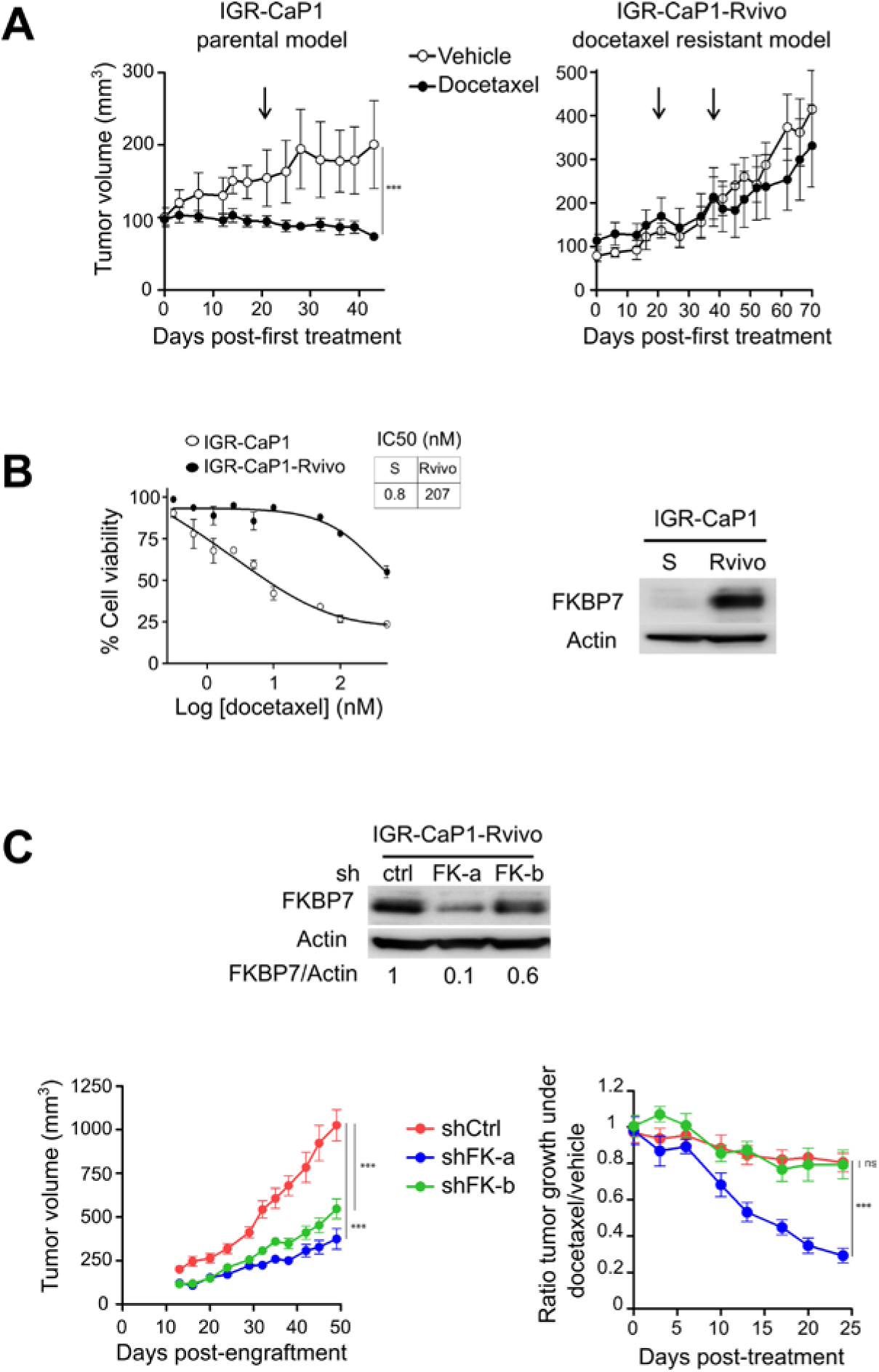
FKBP7 silencing reduces tumor growth in a docetaxel-resistant mouse model (**A**) Tumor growth curves of tumors issued from IGR-CaP1 and IGR-CaP1-Rvivo subcutaneous xenografts after treatment with either vehicle (5% glucose solution) or docetaxel (30mg/kg I.P). Data represent the mean ± SD. N=5 mice per group. Arrows indicate the time of the subsequent docetaxel injections. ***p<0.001 (two-way ANOVA with Bonferroni posttests). (**B**) Left panel: Proliferation assay, determination of IC50 of docetaxel treatment in parental IGR-CaP1 and IGR-CaP-Rvivo cells. The data are represented as the mean ± SD. Cell viability is shown relative to control treatment. Right panel: Immunoblot of FKBP7 in IGR-CaP1 (S) and IGR-CaP1-Rvivo cells. Actin is the loading control. (**C**) Immunoblot shows FKBP7 knockdown efficiency after transduction of the IGR-CaP1-Rvivo cells with lentivirus expressing two shRNAs targeting FKBP7 as compared to control shRNA. Actin is the loading control. Quantification of sh-FKBP7s relative to sh-Ctrl transduced cells was performed with the Image Lab software. Average tumor volume ± SEM obtained from xenografts of IGR-CaP1-Rvivo transduced with control shRNA (n=6) or FKBP7-directed shRNAs (n=7/group) were represented before treatment (left panel). When tumors reached 450-500mm^3^, mice received vehicle or docetaxel (right panel). The ratio between the tumor volume of mice receiving docetaxel and mice receiving vehicle (at days 0 and 21) is represented. shCtrl: n=6/group; shFKBP7s: n=7/group.

### FKBP7 silencing does not affect cell growth in non-tumorous cells

We determined the levels of FKBP7 in various normal human cell lines of different origins (fibroblastic, myoblastic, epithelial, or endothelial cells). These non-cancerous cells exhibited a high protein level of FKBP7, which was even higher than the one observed in IGR-CaP1-Dtx-R cells (Fig S4A). However, contrasting with the results obtained in chemoresistant cells, we observed no effect of FKBP7 inhibition on cell growth and viability in non-cancerous cells (Fig S4B), suggesting that FKBP7 could have a different function in normal and in cancerous cells.

### FKBP7 interacts with eukaryotic translation initiation factors to regulate protein translation

To identify the molecular pathway in which FKBP7 exerts its differential function in non-cancerous and in chemoresistant cells, we performed qualitative and quantitative mass spectrometry analyses. The protein interactome of FKBP7 was identified by performing an immunoprecipitation of endogenous FKBP7 with two different antibodies in both the non-cancerous RPE-1 and in the docetaxel-resistant IGR-CaP1 cell lines (Table S3). Protein network analysis through Ingenuity Pathway Analysis (IPA) indicated that in both cell lines, FKBP7 protein interactors were mainly distributed in the same three intracellular pathways that are associated with protein translation. Specifically, eIF2, eIF4 and mTOR signaling were the most represented pathways in the signatures (Fig 4A). In particular, many eukaryotic translation initiation factors were represented. To elucidate the molecular mechanisms by which FKBP7 depletion exerts its cytotoxic effects in chemoresistant cells only, we used SILAC (stable isotope labeling by amino acids in cell culture) in FKBP7-silenced RPE-1 and IGR-CaP1-Dtx-R cells or in cells transfected with control siRNA. We obtained a list of 910 and 1223 differentially expressed proteins in RPE-1 and in IGR-CaP1-Dtx-R cells, respectively (Table S4). Consistently with previous results, IPA analysis showed that the major pathways affected by FKBP7 silencing were the same three pathways implicated in the control of protein translation. Interestingly, based on the calculated Z-score, these pathways were downregulated in IGR-CaP1-Dtx-R, while they were mainly upregulated in the non-cancerous RPE-1 cells (Fig 4B-C).

**Figure 4.**
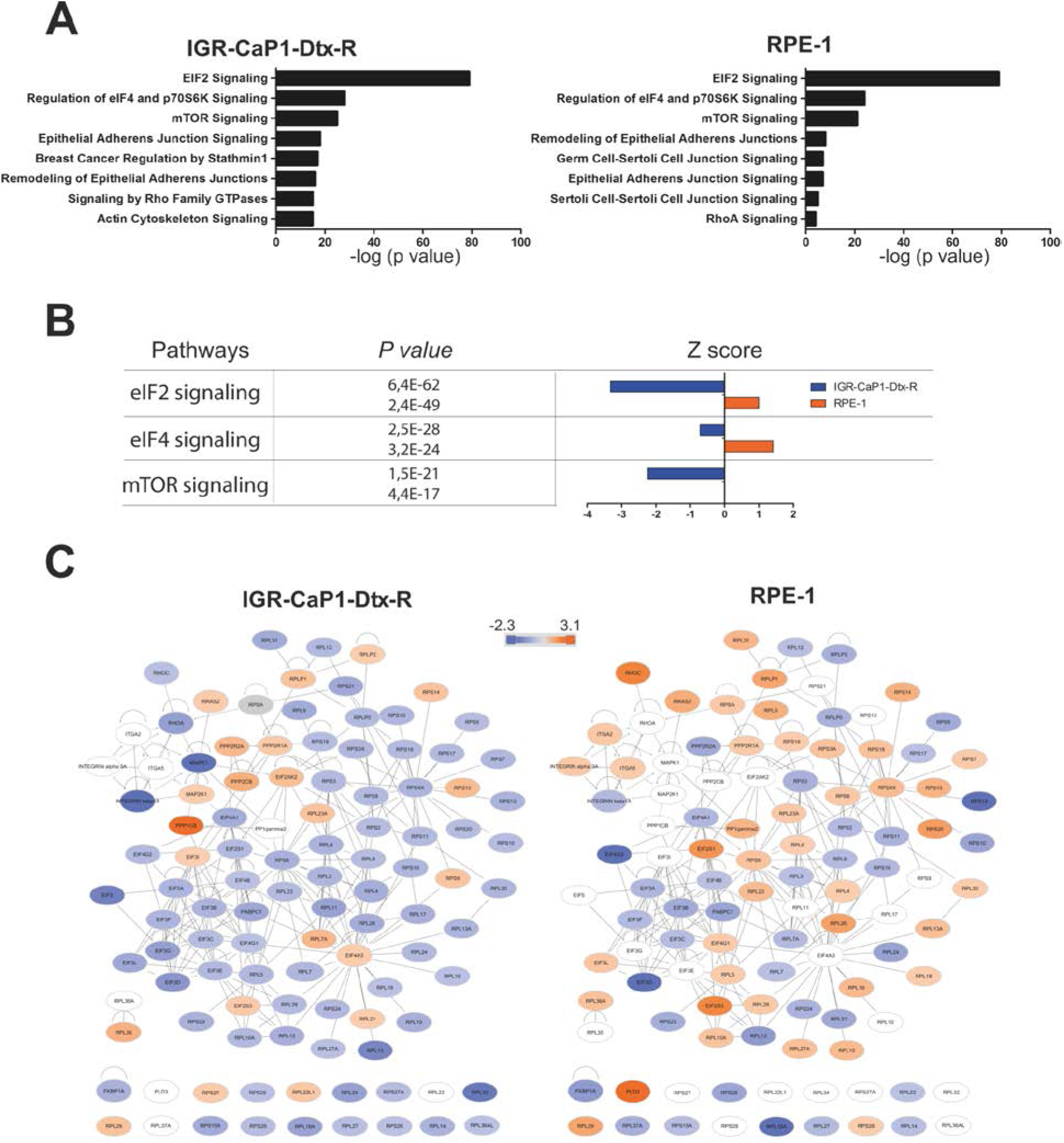
FKBP7 interacts with eukaryotic translation initiation factors to regulate protein translation. (**A**)Ingenuity pathway analysis showing the canonical pathways identified with the specific protein partners of FKBP7 in IGR-CaP1-Dtx-R and RPE-1 from global proteomic IP. (**B**) Major three pathways deregulated in cells transfected with a siRNA targeting FKBP7 (siFK-2), identified with IPA analysis from SILAC data. Each pathway is associated with a p value and a Z-score for IGR-CaP1-Dtx-R and RPE-1. (**C**) Network showing the protein members of eIF2, eIF4 and mTOR pathways deregulated by FKBP7 knockdown in IGR-CaP1-Dtx-R and RPE-1 cells. The color represents the fold change of protein expression between siFKBP7 and siNT.

### FKBP7 regulates the formation of eIF4F translation initiation complex through a direct interaction with eiF4G

All three individual components of the eIF4F cap-dependent mRNA translation complex, eIF4E, eIF4A and eIF4G, were identified as protein partners of FKBP7 in resistant and in RPE-1 cells. EiF4F is an interesting target, known to be involved in the resistance mechanism to many cancer therapies (Cencic *et al*, 2011; Boussemart *et al*, 2014; Robert *et al*, 2014; Zindy *et al*, 2011). We focused on eIF4G, the most downregulated eukaryotic translation factor in IGR-CaP1-Dtx-R but not in RPE-1. FKBP7-silencing led to a decrease in eIF4G in docetaxel-resistant IGR-CaP1 while it led to an increase in eIF4G in the corresponding non-cancerous RPE-1 cells (Fig 5A). In contrast, the protein level of eIF4A and eIF4E remained unchanged upon FKBP7 silencing (Fig 5A). Of note, the level of eIF4G was also lower in cabazitaxel-resistant IGR-CaP1 model and in the chemoresistant 22RV1 cellular model (Fig 5B).

**Figure 5.**
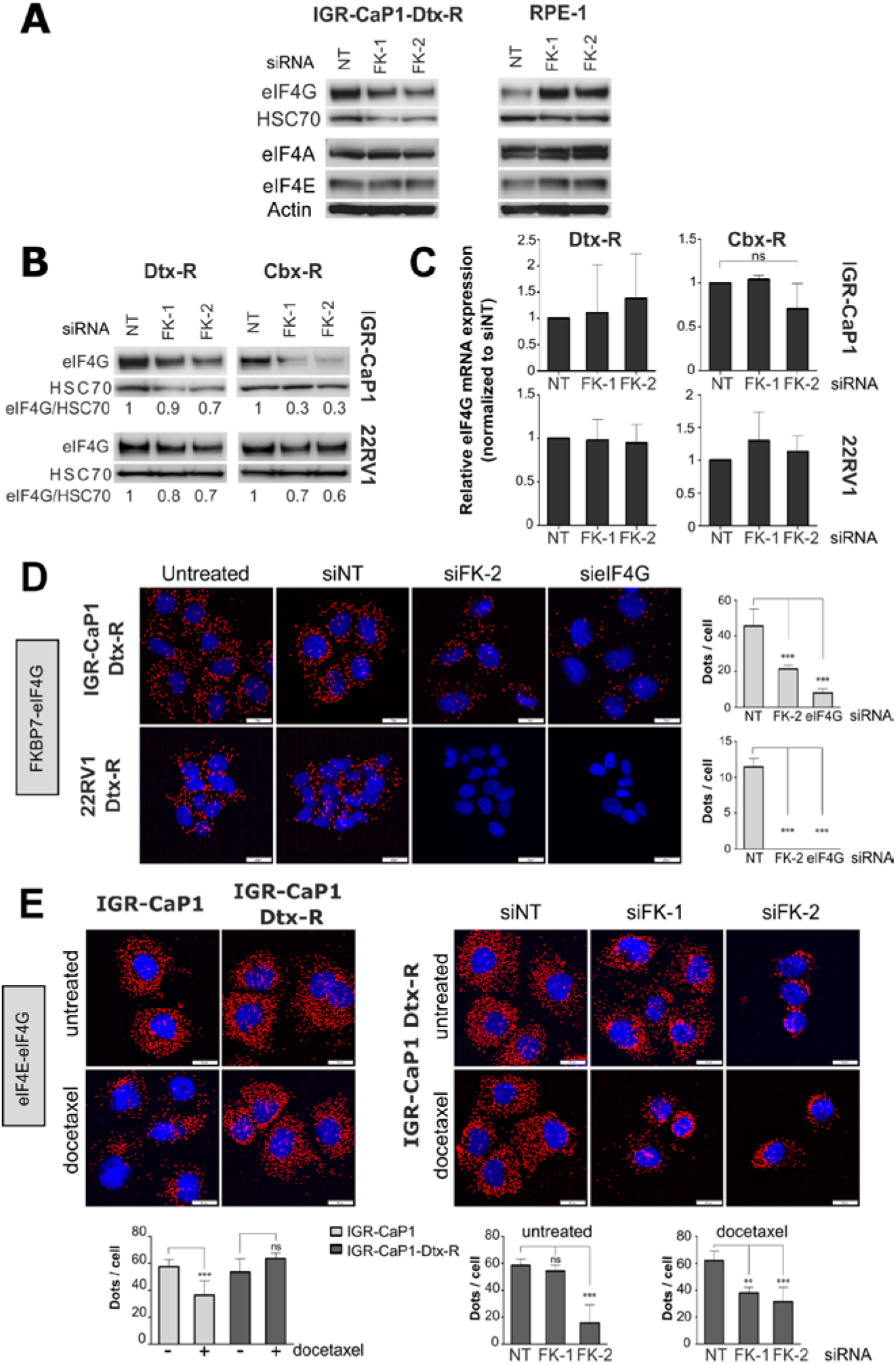
FKBP7 regulates the formation of eIF4F translation initiation complex through a direct interaction with eiF4G. (**A**) Immunoblots showing eIF4G, eIF4A and eIF4E expression in IGR-CaP1-Dtx-R and RPE-1 cells transfected either with siRNA control (siNT) or with two different siRNAs targeting FKBP7 (siFK-1 or siFK-2). HSC70 is the loading control for eIF4G expression and Actin for eIF4A and eIF4E. (**B**) Immunoblots showing eIF4G expression in docetaxel- and cabazitaxel-resistant cells IGR-CaP1 and 22RV1 (Dtx-R and Cbx-R respectively) transfected either with siRNA control (siNT) or with two different siRNAs targeting FKBP7 (siFK-1 or siFK-2). HSC70 is the loading control. The ratio eIF4G/HSC70 was calculated with the Image Lab software. (**C**) Quantitative RT-PCR (qRT-PCR) of eIF4G mRNA in docetaxel- and cabazitaxel-resistant cells IGR-CaP1 and 22RV1 (Dtx-R and Cbx-R respectively) transfected either with siRNA control (siNT) or with two different siRNAs targeting FKBP7 (siFK-1 or siFK-2). EIF4G level was normalized to siNT condition and with GAPDH. (**D**) FKBP7-eIF4G interaction detected by PLA in docetaxel-resistant IGR-CaP1 cells (IGR-CaP1-Dtx-R) and 22RV1 cells (22RV1-Dtx-R). Cells were either untreated or transfected with siRNA control (siNT) or siRNA targeting FKBP7 (siFK-2) or siRNA targeting eIF4G (sieIF4G). Scale bars: 20μm. The interactions were visualized as red dots and the nuclei in blue. Interaction dots were quantified (n≥ 100 cells) and analyzed with the general linear model, *** p<0.001. (**E**) eIF4E-eIF4G interaction detected by PLA in parental and docetaxel resistant IGR-CaP1 cells (left panel). Cells were either untreated or treated with 5nM of docetaxel for 24h. On right panel, eIF4E-eIF4G interactions were detected on docetaxel-resistant IGR-CaP1 cells, either transfected with siRNA control (siNT) or siRNAs against FKBP7 (siFK-1 and siFK-2) alone or combined with 5nM of docetaxel for 24h. Scale bars: 20μm. The interactions were visualized as red dots and the nuclei in blue. Interaction dots were quantified (n≥ 100 cells) and analyzed with the general linear model or with Wilcoxon rank test, ** p<0.01;*** p<0.001.

To understand how FKBP7 silencing could decrease the level of eIF4G, we first analyze the effect of FKBP7 depletion on eIF4G gene expression. As shown in Fig 5C and in Fig S5A, efficient silencing of FKBP7 did not affect the level of eIF4G mRNA in FKBP7-silenced chemoresistant IGR-CaP1 and 22RV1 cell lines. Therefore, our results showed that while eIF4G is still transcribed, its protein level is decreased when FKBP7 is silenced, thus leading to the hypothesis of a regulation of FKBP7 on eIF4G expression at mRNA translation or protein stability levels. We thus investigated the ability of FKBP7 to interact with eIF4G by performing proximity ligation assays (PLA), allowing the interaction to be quantitatively visualized by fluorescent dots. Our results reveal an interaction between FKBP7 and eIF4G in docetaxel-resistant IGR-CaP1 and 22RV1 cells (Fig 5D). This interaction was largely affected in FKBP7- or eIF4G-depleted cells, as shown by the loss in the number of dots per cell (43% and 80% reduction of dots in IGR-CaP1-Dtx-R cells, 93% and 86% in 22RV1-Dtx-R cells, for FKBP7- and eIF4G-silencing respectively) (Fig 5D). In this experiment, the silencing of eIF4G was very efficient (Fig S5B) and the specificity of the FKBP7-eIF4G interaction was validated (Fig S5C).

Since FKBP7 interacts with eIF4G and modulates eIF4G protein levels, we next investigated the effect of FKBP7 on the formation of the eIF4F complex using PLA. Docetaxel treatment led to a decrease in the formation of the eIF4E-eIF4G complex in the docetaxel-sensitive IGR-CaP1 cells (Fig 5E, left panel). Strikingly, but consistently with other studies showing that the formation of the eIF4E-eIF4G determines the sensitivity to anti-cancer targeted therapies (Boussemart *et al*, 2014), this effect was not observed in the resistant cell line IGR-CaP1-Dtx-R (Fig 5E, left panel). SiRNA-mediated depletion of FKBP7 induced a decrease in the formation of the eIF4E–eIF4G complex upon treatment with docetaxel in resistant cells (Fig 5E, right panel). Thus, the formation of the eIF4E-eIF4G complex can be inhibited by docetaxel in resistant cells only when FKBP7 is silenced. Therefore, FKBP7 which is up-regulated in chemoresistant cells could maintain the eIF4F complex by directly regulating eIF4G protein levels and consequently increasing its activity, leading to a hyper-activation of cap-dependent translation and subsequent cell survival upon taxane treatment. Finally, FKBP7 was able to stabilize the eIF4F complex formation upon taxane treatment and thus could be a novel regulator of eIF4F.

### Targeting of chemoresistant prostate cancer cells with small molecule inhibitors of eIF4A

We looked at the structure and the function of FKBP7 to determine whether available drugs might target the chemoresistant cells. FKBP12, which is the target of anti-mTOR rapamycin, shares sequence homology with FKBP7 (Blackburn & Walkinshaw, 2011) and since rapamycin is expected to dephosphorylate 4EBP1, it should disrupt the eIF4F complex. However, we observed that IGR-CaP1-Dtx-R cells were resistant to rapamycin compared to the breast cancer cell line MCF7 (Fig 6A). Since FKBP7 regulates eIF4F complex formation which is involved in chemoresistance, we tested inhibitors of eIF4F on two chemoresistant cell lines with different expression of the androgen receptor. We used three small molecules (silvestrol, FL3, FL23) which were reported to target the eIF4F complex by inhibiting the helicase eIF4A (Bordeleau *et al*, 2008; Cencic *et al*, 2009; Boussemart *et al*, 2014; Chambers *et al*, 2013). Silvestrol induced a cytotoxic effect in the A375 melanoma cell line (Boussemart *et al*, 2014), but showed no effect on the IGR-CaP1-Dtx-R cells (Fig 6A). This lack of activity of silvestrol could be due to the fact that it is a substrate of the Pgp drug efflux pump (Gupta *et al*, 2011) which is highly expressed in the IGR-CaP1-Dtx-R cells (Fig S6). In contrast, the docetaxel-resistant IGR-CaP1 and 22RV1 cells were sensitive to low doses of FL3 (IC_50_=19nM and 14nM respectively) and FL23 (IC50=7nM and 7nM) (Fig 6B). Both compounds target eIF4A but are not substrates of Pgp (Thuaud *et al*, 2009). The observed IC_50_ for these compounds were similar to the ones observed in the melanoma A375 cells (IC_50_=13 and 18nM respectively). Thus, our results showed that FL3 and FL23 were able to kill the chemoresistant cells; however, no synergy was observed when these translation inhibitors were used in combination with docetaxel (Fig S7A). We also confirmed the cytotoxic effect of these two flavaglines on parental, docetaxel- and cabazitaxel-resistant cell lines, as shown by the induction of PARP cleavage (Fig 6C). Importantly, FL3 and FL23 did not lead to apoptosis of RPE-1 cells (Fig 6C), reinforcing the interest of targeting eIF4A to abrogate taxane resistance in prostate cancer.

**Figure 6.**
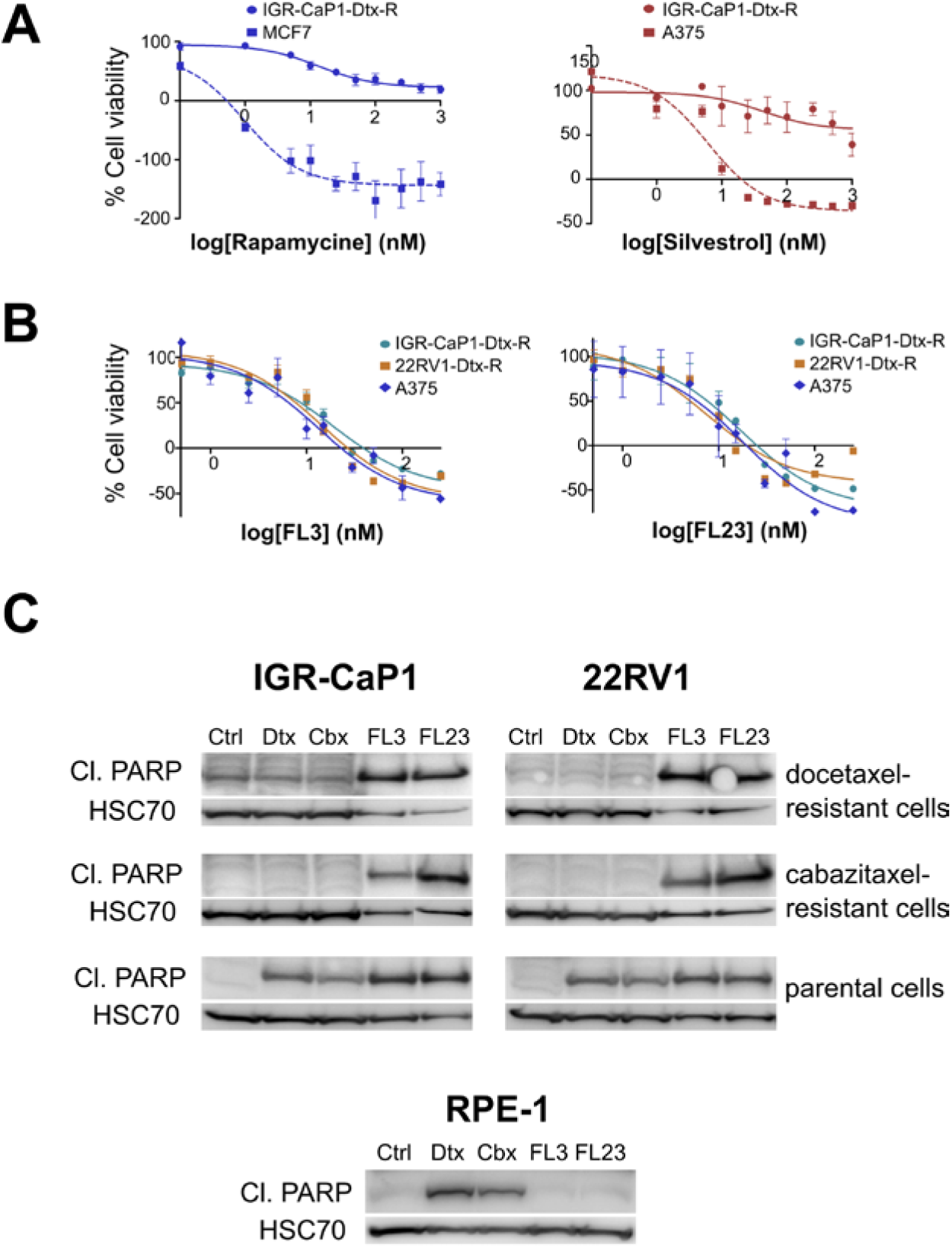
Targeting of chemoresistant prostate cancer cells with small molecules inhibitors of eIF4A. (**A**) Proliferation assay, cell viability (calculated relatively to control treatment) of IGR-CaP1-Dtx-R and the breast cancer MCF7 cells after 48h of treatment with rapamycin (left panel), of IGR-CaP1-Dtx-R and the melanoma A375 cells after 48h of treatment with silvestrol (right panel). The data are represented as the mean ± SD. (**B**) Proliferation assay, cell viability (calculated relatively to control treatment) of docetaxel-resistant cells IGR-CaP1 and 22RV1 (IGR-CaP1-Dtx-R and 22RV1-Dtx-R respectively) and the melanoma A375 after 48h of treatment with either flavagline 3 (FL3, left panel) or flavagline 23 (FL23, right panel). The data are represented as the mean ± SD. (**C**) Immunoblots of cleaved (Cl.) PARP protein in parental, docetaxel- and cabazitaxel-resistant IGR-CaP1 and 22RV1 cells and in RPE-1 cells, either untreated (ctrl), treated for 72h with 10nM docetaxel (Dtx) or 5nM cabazitaxel (Cbx) or treated for 48h with 150nM flavagline FL3 or FL23. HSC70 is the loading control.

## DISCUSSION

Resistance to chemotherapy represents a major challenge for cancer therapy (Rebucci & Michiels, 2013; Zhang *et al*, 2015). Understanding the mechanisms of chemoresistance and identifying biomarkers for chemoresistance is crucial for developing new therapeutic strategies and overcoming drug resistance. In this study, we make the unprecedented observation that FKBP7 is an important determinant of prostate cancer cell response to taxanes. Indeed, acquisition of resistance to taxane upon long term-treatment of prostate cancer cells with docetaxel or cabazitaxel is correlated with increased levels of FKBP7. In addition, the identification of the FKBP7 signaling network allowed us to link the participation of FKBP7 in chemoresistance to the initiation step of protein translation and more precisely, the eIF4F translation initiation complex. Using eIF4A inhibitors at nanomolar range concentrations allowed us to circumvent docetaxel resistance and cabazitaxel resistance in two prostate cancer chemoresistant cell lines representing the most challenging models to sensitize to taxanes.

Recently, several FKBP proteins have been shown to participate in cancer progression and chemoresistance (Solassol *et al*, 2011). FKBP5 is known to regulate steroid receptor activation and prostate cancer progression (Storer *et al*, 2011). It was previously shown that FKBP5 expression levels correlate with the sensitivity of tumor cells to chemotherapeutic agents (Li *et al*, 2008). In addition, FKBP5 negatively regulates the Akt kinase by acting as a scaffold protein for Akt and the phosphatase PHLPP and thus increases chemosensitivity (Pei *et al*, 2009). FKBP5 was also shown to be involved in the resistance of malignant melanoma and in the resistance to taxol in ovarian cancer cells (Sun *et al*, 2014). Interestingly, as we observed with FKBP7 (data not shown), FKBP5 overexpression was only observed upon treatment of cells with microtubule-targeting agents and was not observed with DNA-damaging agents such as cisplatin (Sun *et al*, 2014). The specificity of this response to microtubule-targeting agents may be related to the regulatory role of FKBPs on the cytoskeletal proteins observed for FKBP4 and FKBP5 (Cioffi *et al*, 2011). Other FKBPs such as FKBPL were shown to have therapeutic and biomarker potential in cancer (Robson & James, 2012). The implication of endoplasmic reticulum FKBPs in carcinogenesis was only reported for FKBP10 (Paulo *et al*, 2012; Olesen *et al*, 2005; Quinn *et al*, 2013) and recently for FKBP14 (Lu *et al*, 2016; Huang *et al*, 2016).

Our study reveals a functional role of the FKBP7 chaperone in acquisition of docetaxel and cabazitaxel resistance in prostate cancer, which is observed both in AR-positive and in AR-negative cells and is independent of the ABCB1/MDR1 drug efflux pump expression. The fact that FKBP7 is overexpressed in human prostate tumors corroborates our *in vitro* observations, as our clinical data showed that high expression of FKBP7 is more frequent in tumors compared to normal tissues and is correlated with a lower time to recurrence in patients who received taxane as neoadjuvant chemotherapy. A high level of FKBP7 correlates with a bad prognosis, which suggests that FKBP7 expression could provide a relevant tissue marker of taxane-resistance. Pre-clinical evidence in a docetaxel-resistant mouse model confirmed that FKBP7 expression sustained the growth of taxane-resistant prostate cancer cells. Therefore, our results implicate a previously uncharacterized member of the PPIase family in the mechanism of resistance to microtubule-targeting agents.

Although high expression of FKBP7 is observed in non-cancerous and chemoresistant cells, FKBP7 silencing triggers cell death in taxane-resistant tumor cells only. This suggests that survival of resistant cells may be attributed to a novel function of FKBP7 in these cells. The comparison of resistant cells and non-cancerous cells in a large proteomic approach allowed us to identify the eukaryotic translational initiation factor and mTOR pathways as the main regulatory networks linked to FKBP7. These pathways are deregulated in resistant tumor cells compared to normal cells. In addition, we identified the eIF4G component of the eukaryotic initiation factor eIF4F as the major downstream target of FKBP7 and showed that the interaction between FKBP7 and eIF4G controls eiF4G protein level. Translation initiation is a highly regulated biological process that has been hijacked by tumor cells to increase the rate of protein synthesis for specific genes and promote cell survival. EIF4G is a large scaffolding protein which binds the eIF4A helicase and the eIF4E cap-binding protein to form the heterotrimeric eIF4F complex for mRNA translation. Accumulation of evidence showed that increased activity of eIF4F contributes to the malignant transformation process via the increase in translation of a limited set of pro-oncogenic mRNA transcripts (Bhat *et al*, 2015). We show that the FKBP7-eIF4G interaction led to an increased level of eIF4G which is normally a limiting factor of this process. Through its chaperone activity, FKBP7 may protect eIF4G from degradation by preventing its ubiquitin-mediated proteolysis. This function could also be achieved by the Hsp90 and Hsp70 proteins which are only associated with FKBP7 in resistant cells (Table S3). Such a process has been described for Hsp27 and eIF4E (Andrieu *et al*, 2010). Alternatively, as shown for FKBP10 (Stocki *et al*, 2016), conformational changes of the target protein induced by the peptidyl-prolyl isomerase activity of FKBP7 could also be responsible for the stability of eIF4G. Thus, our data suggest that the survival of chemoresistant cells may be attributed to an over-stimulation of eIF4F, one of the downstream effectors of the Akt-mTOR pathway mediated by FKBP7 overexpression (Fig 7).

**Figure 7.**
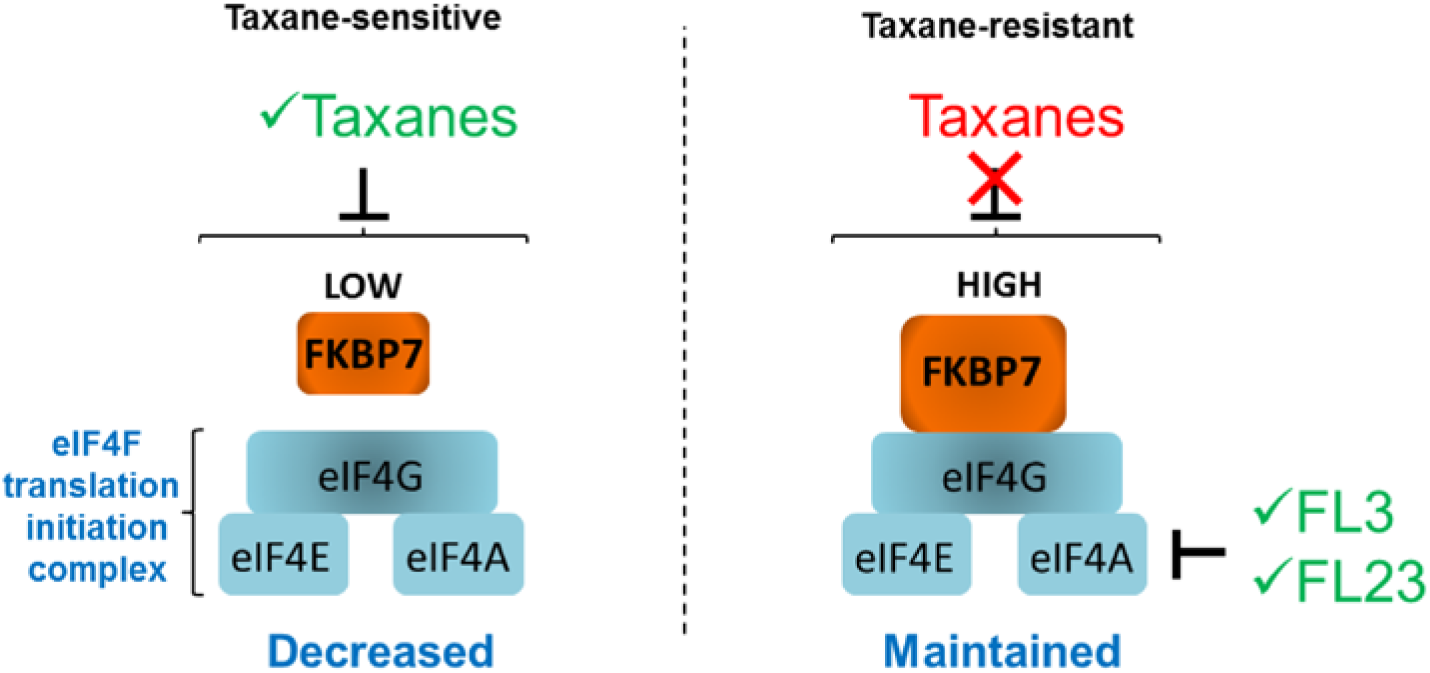
Mechanistic model of FKBP7 role in prostate cancer cells. By directly binding to eIF4G, overexpression of FKBP7 in chemoresistant cells leads to an increased translational activity of eIF4F, and thus to survival of chemoresistant cancer cells. Inhibitors of this pathway are indicated. Only the flavaglines (FL3 and FL23) efficiently inhibited this pathway, leading to cell death of chemoresistant cells.

Hyperactivity of the Akt-mTOR pathway was observed in many cancers including prostate cancer (Bitting & Armstrong, 2013) and this pathway was shown to be implicated in chemoresistance in prostate cancer (Vidal *et al*, 2015). In that case, the chemoresistance should be alleviated with treatment of resistant cells with the mTOR inhibitor rapamycin. However, we observed a modest inhibitory effect of rapamycin in our docetaxel-resistant IGR-CaP1 model. Furthermore, in the clinic, rapamycin and rapamycin-analogues showed only modest effects (Vaishampayan *et al*, 2015; Bitting & Armstrong, 2013). In light of our results, the development of novel strategies targeting FKBP7 or eIF4F would be of particular interest to overcome chemoresistance in prostate cancer.

As we have shown that FKBP7 has an important role in survival networks that protect cancer cells against therapeutic agents, FKBP7 could be an interesting target in prostate cancer. But to date, structural analysis of the full-length FKBP7 protein is not available for the design of specific inhibitory molecules, and although screening strategies have been developed to discover new drugs inhibiting FKBP activity, relatively few compounds have been reported and designing isoform specific inhibitors is still challenging (Blackburn & Walkinshaw, 2011). Thus, the design of small molecule ligands with specificity for FKBP7 will be the focus of our drug research in the very near future. The discovery of the interaction between FKBP7 and eIF4G led us to assess the efficacy of inhibitors, which directly target eIF4F complex. Targeting the translation machinery is a promising strategy to minimize acquired resistance (Bhat *et al*, 2015). We first tested silvestrol which inhibits eiF4A and was shown to provide therapeutic benefits in prostate cancer xenografts (Cencic *et al*, 2009). Unfortunately, silvestrol shows no obvious efficacy towards docetaxel-resistant IGR-CaP1 cell proliferation, which could be attributed to the high expression of ABCB/MDR1 in this model. However, flavaglines FL3 and FL23, which also target eIF4A, are highly cytotoxic in docetaxel-resistant models with IC_50_ in the range of 10nM, independent of drug-efflux pumps. These synthetic compounds (Thuaud *et al*, 2011; Ribeiro *et al*, 2012) were shown to overcome multidrug resistance *in vitro* (Thuaud *et al*, 2009). In addition, FL3 and FL23 have also been recently shown to alleviate resistance to vemurafemib in melanoma (Boussemart *et al*, 2014). Our findings open a new avenue in the field of taxane resistance, which now needs to be explored in other cancer types. We propose that targeting eIF4F is a promising therapeutic strategy to overcome both docetaxel- and cabazitaxel-resistance.

Finally, our study reveals the critical role played by the chaperone FKBP7 in acquired taxane-resistance in prostate cancer and its potential for development as a predictor of chemoresistance. We also found a novel therapeutic strategy to overcome taxane resistance by targeting the eIF4F translation initiation complex.

## METHODS

### Cell culture and reagents

The parental and taxane-resistant IGR-CaP1, PC3, LNCaP and 22RV1 human PCa cell lines were maintained in RPMI1640 medium supplemented with 10% FBS. Taxane-resistant clones were selected by exposing cells to docetaxel or cabazitaxel in a dose-escalation manner as described for IGR-CaP1 (Nakouzi *et al*, 2014). Resistant cell lines were treated monthly with the maximal dose of docetaxel or cabazitaxel to maintain the resistant phenotype. The non-tumorigenic prostate epithelial cells RWPE-1 were maintained in Keratinocyte Serum Free Medium (K-SFM) (Thermo Fisher Scientific), supplemented with bovine pituitary extract and human recombinant epidermal growth factor (Thermo Fisher Scientific), the Retina Pigmented Epithelium-1 (RPE-1) cells were maintained in DMEM:F12 (GIBCO) with 10% FBS (Thermo Fisher Scientific), the Human Kidney-2 cells (HK-2) and the Human Umbilical Vein Endothelial cells (HUVEC) were maintained in DMEM (GIBCO) with 10% FBS, the fibroblasts from lung (MRC-5) were maintained in EMEM (GIBCO) with 10% FBS and the Human immortalized myoblasts (LHCN-M2) were maintained in DMEM (GIBCO) and medium 199 (Invitrogen) with 20% FBS. Docetaxel was purchased from Sanofi-Aventis and cabazitaxel from Selleckchem.

### Microarray

Gene expression was profiled using a 4×44K Human Whole Genome (G4112F) expression array (Agilent Technologies) with a dual-color dye-swap competitive hybridization procedure, according to the manufacturer’s instructions. Total RNA from untreated parental IGR-CaP1 cells was used as the RNA reference. Total RNA from IGR-CaP1 cells resistant to 5nM, 12nM, 25nM 50nM, 100nM, and 200nM of docetaxel respectively, were used as samples. Two independent replicates of each sample were used. Image analyses (quantification, normalization) were performed with Feature Extraction software (Agilent Technologies) and gene expression analysis was performed using Bioconductor. Analysis of genes differentially expressed between parental and resistant cell lines was performed as follow.

For all the resistant cell lines, Log10 (ratios) were computed against the sensitive cell line. In order to select relevant genes, we combined three different strategies. First, gene permanently overexpressed (or under-expressed), from the first dose, in all the resistant cell lines, were tested using multiple t-tests with boostraps consisting in 10,000 resamples with replacements. The resulting p-values where adjusted with Benjamini-Hochberg correction method (Giraldo *et al*, 2002). Adjusted p-values ≤0.05 were declared as significant. Secondly, genes with monotonic increasing (or decreasing) expression over the increasing doses were tested using 5-parameter logistic regressions (Giraldo *et al*, 2002). The decision rule combined an absolute fold change of at least 2 between the lower and the upper asymptotes, and an adjusted p-value ≤1e-3, representing the quality of the correlation between the fitted and observed values. Eventually, supplementary potentially informative genes were selected using an information criterion method (ongoing publication). Briefly, this method uses a reversed principal components analysis, where probes are considered as observations. For each gene, an information criterion is computed in order to quantify its ability to separate samples. This analysis allowed the elaboration of 998 genes potentially implicated in docetaxel resistance comprising 593 upregulated genes and 407 down-regulated genes.

### Image-based high-content siRNA screening

IGR-CaP1-R100 cells were treated with 100nM of docetaxel for three weeks after thawing. Cells were then plated in 384-well plates (ViewPlate-384 Black optical clear bottom, Perkin Elmer) at 1,500 cells/well and were allowed to adhere overnight. A library of siRNAs (Custom FlexiPlate Qiagen set) targeting 593 different genes (4 siRNAs per gene) corresponding to overexpressed genes of the gene signature were transfected at 10nM using 0.05 μl Interferin (Polyplus Transfection) in 60μl total volume. siKIF11, siGL2 and siGOLGA2 siRNAs were used as controls. Forty-eight hours after transfection, cells were treated or not with 100nM docetaxel. The effect of siRNAs was addressed by high-content immunofluorescence imaging. After 72 hours, cells were fixed with 4% (w/v) formaldehyde for 15min and washed with phosphate buffered saline (PBS). Cells were next quenched with 0.05M NH4Cl and permeabilized first with a PBS solution containing 0.2% BSA and 0.05% saponin, and subsequently with 0.5% Triton-X100. Cells were then incubated for 60 min with mouse anti-Ki67 (1:500; Millipore) and EdU (10 μM; Invitrogen). Next, cells were washed twice and incubated with Alexa Fluor 488-coupled secondary antibodies (1:400; Jackson ImmunoResearch). Cell numeration was determined by 0.2μg/ml DAPI staining. The screening was realized in triplicate. Images were acquired with an INCell 2000 automated wide-field system (GE Healthcare). The anti-proliferative effect of each siRNA-treated well was evaluated using dual-parameter measurements of nuclei staining fluorescence in Ki-67 and EdU channels. Images were quantified in each well with the INCell Analyzer workstation software (GE Healthcare).

### Data analysis and hit calling

Data correction and scoring were performed as described previously (Mahmood *et al*, 2014). Briefly, data were first transformed with log or logit functions. B-score normalization was then applied to each replicate, separately, and included corrections for plates, rows and columns (Brideau *et al*, 2003; Malo *et al*, 2006). Median and median absolute deviation (MAD) were computed and used to compute Robust Z-scores (RZ-scores) for each sample, according to the formula: score = (value - median)/(1.4826x Median MAD) (Birmingham *et al*, 2009). RZ-scores were calculated for the comparison of each siRNA against the GL2 negative control population. A gene was identified as a ‘hit’, if the RZ-score for at least two of four siRNAs was > 2 or < -2 in at least two of three replicates. Thirty-four hits were thus selected.

### Immunohistochemistry

The prostate tissue samples used for TMA were obtained from the Vancouver Prostate Centre Tissue Bank with written informed patients’ consent, clinical information and institutional study approval, and were previously reported. The H&E slides were reviewed and the desired areas were marked on them and their correspondent paraffin blocks. For the first study, 8 TMAs were manually constructed (Beecher Instruments) by punching duplicate cores of 1mm for each sample. All specimens (n=381) were obtained through radical prostatectomy, except CRPC samples that were obtained via transurethral resection of the prostate. For the second study, 2 TMAs were manually constructed from 90 patients who received docetaxel with radical prostatectomy. Analysis was performed on 69 selected patients with presence of cancer and good core integrity. Immunohistochemical staining with FKBP7 antibody (Sigma-Aldrich) was conducted by Ventana autostainer model Discover XT™ (Ventana Medical System) with enzyme labeled biotin streptavidin system and solvent resistant DAB Map kit. All stained slides were digitalized with the SL801 autoloader and Leica SCN400 scanning system (Leica Microsystems) at a magnification equivalent to ×20. The images were subsequently stored in the SlidePath digital imaging hub (DIH; Leica Microsystems) of the Vancouver Prostate Centre. Representative cores (clearly positive, clearly negative and mixed positive/negative) were manually identified by the pathologist. Values on a four-point scale were assigned to each immunostain. Descriptively, 0 represents no staining by any tumor cells, 1 represents a faint or focal, questionably present stain, 2 and 3 represents a stain of convincing intensity in a majority of cells.

### Clonogenicity assay

Parental and chemoresistant cells were plated in 6-well plates at low density. Cells were treated with either docetaxel or cabazitaxel for 3 days. After 2 weeks, clones were methanol fixed and stained using Crystal violet (Sigma-Aldrich). Clones were counted manually.

### SiRNA transfection

Cells were plated in 96-well plates with 20nM siRNA either control, siNT (Thermo Scientific) or targeting FKBP7 gene (Stealth RNAiTM, siFKBP7-1: HSS122495 and siFKBP7-2: HSS182176). Reverse transfection was performed using lipofectamine RNAiMax (Thermo Fisher Scientific). Cell viability was assessed every day for 4 days with WST-1 Cell Proliferation Reagent (Roche). The transfection efficiency was checked by western-blot. Cells were plated in 6 well-plates and 48 hours after transfection with siRNAs, whole cell extracts were prepared in RIPA buffer complemented with proteases inhibitors (Roche) and phosphatase inhibitors (Sigma).

### Establishment of the in vivo docetaxel resistant model

All animal experiments were approved by the local ethics committee (CEEA IRCIV/IGR no. 1226.01, registered with the French Ministry of Research) and were in compliance with EU Directive 63/2010. IGR Animal Resources holds a National Institutes of Health; Department of Health and Human Services Animal Welfare Insurance (no. A5660-01) and were in compliance with the Guide for the Care and Use of Laboratory Animals. Six-week-old male athymic nude mice (NC-nu/nu) were purchased from the animal facility of Gustave Roussy (GR). Ten millions of IGR-CaP1 cells resistant to 12nM, 25nM and 50nM were implanted subcutaneously in the right rear flank region. Tumor growth was monitored using a digital caliper. Tumors volumes were calculated with the ellipsoid volume formula Lxl2x0.5, where L is the length and l the width. After six months of follow-up, only one tumor issued from the IGR-CaP1-R25 xenograft was measurable and subsequently transplanted into other nude mice. Five successive transplantations of this tumor were performed in mice, which were injected I.P. with a single dose of docetaxel, with increasing doses between the passages (starting from 10mg/kg). The resistance model was established after the firth passage. Tumors were excised, digested with enzymes, and the cell suspension was seeded in plastic dishes. The cell line obtained was named IGR-CaP1-Rvivo. For the validation of the resistant model, six-week old male athymic mice were injected S.C. with either 2×10^6^ IGR-CaP1-Rvivo cells or 10×10^6^ IGR-CaP1 cells in 100μl PBS and 50% Matrigel (Corning). When tumors reached an average volume of 100mm^3^, equivalent group of mice (n=5) were injected intraperitoneally with docetaxel (30mg/kg, 3 times) or vehicle every 3 weeks. Tumor growth was monitored weekly during 50-70 weeks using a digital caliper. The mice were sacrificed, and tumors were excised and measured.

### In vivo shRNA knockdown

Two distinct shRNAs targeting FKBP7 (shFK-1 and shFK-2) were engineered and packaged using the lentivirus delivery system. The shRNA plasmids targeting FKBP7 (shFK-1: V3LHS_320392 and shFK-2: V2LHS_270816) and the control plasmid were obtained from Thermo Scientific (GIPZ Lentiviral shRNA). Lentivirus that expressed shRNA was produced using HEK293T cells with the packaging system, pSPAX2 (Addgene plasmid 12260) and pMD2.G (Addgene plasmid 12259). 2×10^6^ IGR-Rvivo cells, transduced with either sh-ctrl or with the two sh-FKBP7s were then injected S.C. in 100μl PBS with 50% matrigel in NSG (NOD Scid gamma) mice purchased from the animal facility of GR. Tumor growth and tumor volume were monitored as described before. When tumors reached an average volume of 450-500mm^3^, mice were injected I.P. with docetaxel or vehicle (30mg/kg) three times, every 3 weeks.

### Immunoprecipitation of endogenous FKBP7

IGR-CaP1-Dtx-R and RPE-1 cells were grown in 4 T150 flasks and harvested at ~80% confluency. Cells were lysed in an optimized buffer (120mM NaCl, 20mM Hepes, 1mM EDTA, 5% glycerol, 0.5% NP40 and protease inhibitor) for 30 minutes. The lysate was centrifuged and the pellet was re-extracted again in the same buffer. The two supernatants were pooled and incubated overnight with 6μg of either control IgG (Thermo Fisher Scientific) or two different FKBP7 antibodies (Thermo Scientific, PA5-30629 and Genetex, clone N1C3, GTX116186) and 50μl of magnetic beads (Dynabeads^®^ Protein A, Thermo Fisher Scientific). Afterwards, the beads were washed four times and the protein complexes were eluted twice in LDS sample buffer containing sample reducing agent (Thermo Fisher Scientific). The sample was loaded on NuPage 10% Bis-Tris protein gel; the proteins were allowed to enter the gel after applying 150V voltage for 5 minutes. The band containing the proteins was excised and processed according to the standard protocol (Yoav Benjamini 1995). The analyses of peptide mixtures were performed on EASY 1000nLC + Q-EXACTIVE (Thermo Fisher Scientific), using EASY-Spray nanocolumn (ES800 15cm 75um), 300 nl/min flow and 2h gradient of acetonitrile + 0.1% Formic acid (starting concentration 5% Acetonitrile and final 35%). The mass resolution for the full scan was set at 70000 at 400 m/Z. The 10 most intense precursor ions from a survey scan were selected for MS/MS fragmentation using HCD fragmentation with 27% normalized collision energy, and were detected at a mass resolution of 17,500 at 400 m/z. Dynamic exclusion was set for 30 seconds with a 10 ppm mass window. Each sample was analyzed in triplicates. The acquired data were analyzed with Proteome Discoverer Software package using Mascot search engine. MS/MS spectra were searched with a precursor mass tolerance of 10 ppm and fragment mass tolerance of 0.05 Da. Trypsin was specified as protease with maximum two missed cleavages allowed. The minimal peptide length was specified to be 6 amino acids. The data were searched against a decoy database, and the false discovery rate was set to 1% at the peptide level. FKBP7 interactors were selected when proteins were only identified in the FKBP7 immunoprecipitation as compared to IgG control or when proteins were enriched at least three fold in the specific immunoprecipitation as compared to the control. The specific protein interactors were processed through Ingenuity Pathway Analysis^®^ (IPA).

### SILAC analysis

IGR-CaP1-Dtx-R or RPE-1 cells were respectively adapted to RPMI or DMEM:F12 SILAC media containing either 12C6, 14N4 L-Arginine-HCl + 12C6 L-Lysine-2HCl (light media) or 13C6, 15N4 L-Arginine-HCl + 13C6 L-Lysine-2HCl (heavy media) (Thermo Fisher Scientific) for a minimum of 5 cell doublings. Cells grown in heavy media were transfected with siRNA ctrl by reverse transfection and the cells cultured in light media were transfected with siRNA against FKBP7 (HSS182176). Equal amounts of cells were mixed and lysed in LDS sample buffer and reducing agent (Thermo Fisher Scientific). The sample was loaded on NuPage 10% Bis-Tris protein gel; the proteins were allowed to enter the gel after applying voltage 150V for 5 minutes. The band containing the proteins was excised and processed according to the standard protocol (Shevchenko *et al*, 2006).

The analysis of the obtained peptide mixtures was performed as described above, except the H/L ratio of for each peptide was calculated by the quantitation node. The average of the ratio Light/Heavy was calculated for each protein identified and the lists of proteins in IGR-CaP1-Dtx-R and RPE-1 were processed through IPA.

### Quantitative real-time RT-PCR

TaqMan gene expression assays were used for human *FKBP7* gene (Hs00535040_m1, Thermo Scientific), with *GUSB* as endogenous control (4333767, Applied Biosystems). Syber gene expression assay was also used for human *EIF4G* gene (forward: TGTGGGACTCTTCAGTGCAA, reverse: TGGGATTCTGAAGGGCTATG) with *GAPDH* as endogenous control (forward: GAAGGTGAAGGTCGGAGTC, reverse: GAAGATGGTGATGGGATTTC). Real-time quantification was performed using either TaqMan gene expression master mix or Power SYBR^®^ Green PCR Master Mix on Viia7 System (Applied Biosystems). The ΔΔCT method was used to quantify transcripts.

### Western Blot analysis

Immunoblots were performed from whole cell lysate prepared with RIPA buffer complemented with proteases inhibitors (Roche) and phosphatase inhibitors (Sigma-Aldrich). The following antibodies were used: FKBP7 (1:500, Sigma-Aldrich), PARP (1:1000, Cell signaling), Mdr-1 (1:1000, Santa Cruz biotechnology), eiF4A, eiF4E and eiF4G (2490, 2067 and 2498 respectively, Cell Signaling). β-actin (Sigma-Aldrich) and HSC70 (Santa-Cruz Biotechnology) were used as a loading control. Immunoblot analyses were performed using the enhanced chemoluminescence-based detection kit (Pierce).

### Proximity ligation assay

Interactions between eIF4G and FKBP7 (FKBP7-eIF4G) and eiF4E and eiF4G (eiF4E-eiF4G) were detected by in situ proximity ligation assay (PLA) (Duolink, Sigma-Aldrich). Cells were fixed with paraformaldehyde, permeabilized and the PLA protocol was followed according to the manufacturers’ instructions (Olink Bioscience). After blocking, the primary antibodies were incubated 1h at 37°C. For FKBP7-eiF4G interaction, the antibodies were used at the following concentrations: eiF4G (mouse,Clone A-10, 1:200), FKBP7 (rabbit, Thermo Scientific, PA5-30629, 1:50). For eiF4E-eiF4G: eiF4E (mouse, Clone A-10, Santa Cruz, 1:500), eiF4G (rabbit, 2498, Cell signaling, 1:500). PLA minus and PLA plus probes containing the secondary antibodies conjugated to oligonucleotides were added and incubated for 1h at 37°C. Ligase was used to join the two hybridized oligonucleotides into a closed circle. The DNA was then amplified (with rolling circle amplification), and detection of the amplicons was carried out using the Far red detection kit for fluorescence. Cell nuclei were stained with 4’,6-diamidino-2-phenylindole (DAPI). The slides were mounted with Olink Mounting Medium. Image acquisition was performed with a Virtual Slides microscope VS120-SL (Olympus, Tokyo, Japan): magnification 20X, air objective (0.75 NA), 10-ms exposure for the DAPI channel and 300-ms exposure for the Cy5 channel; 1pixel=0.32μm) and the number of PLA signals per cell was counted using Image Analysis toolbox in Matlab (2011a).

### Statistical Analysis

Statistical analysis was carried out with GraphPad software or by on-site bio-statisticians. For the tissue macro-array experiment, FKBP7 staining quantification in prostate tissues was analyzed with Chi-square test and the percentage of recurrence was calculated with a Fisher test. Survival analysis was performed using the Kaplan-Meier method and curves were compared with the Cox proportional hazard model. Cell proliferation curves of chemoresistant cells transfected with siNT or with siFKBP7s or the different tumor growth curves observed either between vehicle and docetaxel treatment or between sh-ctrl and sh-FKBP7s were analyzed by one two-way ANOVA with Bonferroni posttests. The significance of eIF4G mRNA level with FKBP7 silencing was set with a Paired t-test. The significance of eIF4G-FKBP7 and eIF4E-eIF4G interactions was tested with the general linear model or with Wilcoxon rank test.

## ACCESSION NUMBER

Microarray experiments have been submitted to Array Express data base (EBI, UK) (http://www.ebi.ac.uk/arrayexpress/) with the following accession number E-MTAB-4869.

## ACKNOWLEDGMENTS

We gratefully thank A. Lescure and S. Tessier of the Biophenics staff, the platform of preclinical evaluation AMMICA, the bioinformatic Core Facility, the Development in Pathology Group (UMR981), and S. Shen (UMR981). We thank and pay tribute to Vasily Ogryzko (UMR8126) who brought us a precious help with proteomics but who disappeared before this publication. RWPE-1 was a kind gift from G. Mouchiroud (University Claude Bernard, Lyon), HK-2 from S. Gad-Lapiteau (UMR1186, Villejuif) and HUVEC from S. Rodrigues-Ferreira (UMR981, Villejuif). This work was supported by grants from: INSERM, the Université Paris-Sud11, the grant PAIR PROSTATE program n°2010-1-PRO-03 from the INCA, the ARC Foundation, the Ligue contre le cancer, the ECOS-Sud A10S03 program, AMGEN, the Paris Alliance of Cancer Research Institutes program, “Investissements d’Avenir” an initiative of the French Government and implemented by ANR with the reference ANR-11-PHUC-002, the TA program of GR for genomic and proteomic analyses, and Terry Fox New Frontiers Program Project Grant TFF116129. MGa was supported by the Idex Paris-Saclay fellowship and the ARTP, NJ-PM was supported by the PARRAINAGE CHERCHEUR charity program of GR.

## AUTHOR CONTRIBUTIONS

AC conceived the project; MGa, N J-PM, CG, VO, NAN, LF, EDN, SL, ALP and HA-H performed the experiments and analyzed data; FC performed bioinformatics analysis; JC, FP, MG, YL, SV, LD and KF provided expertise and feedback; MG and LD provided useful reagents; MGa and AC wrote the manuscript; AC and KF secured funding for the project. All authors analyzed the data, discussed results and commented on the manuscript.

